# Revisiting a classical theory of sensory specificity: assessing consistency and stability of thermosensitive spots

**DOI:** 10.1101/2023.07.17.549302

**Authors:** Ivan Ezquerra-Romano, Michael F. Clements, Steven di Costa, Gian Domenico Iannetti, Patrick Haggard

## Abstract

Thermal sensitivity is not uniform across the skin, and is particularly high in small (∼1mm^2^) regions termed ‘thermosensitive spots’. These spots are thought to reflect the anatomical location of specialised thermosensitive nerve endings from single primary afferents. Thermosensitive spots provide foundational support for “labelled line” or specificity theory of sensory perception, which state that different sensory qualities are transmitted by separate and specific neural pathways. This theory predicts a highly stable relation between repetitions of a thermal stimulus and the resulting sensory quality, yet these predictions have rarely been tested systematically. Here we present the qualitative, spatial and repeatability properties of 334 thermosensitive spots on the dorsal forearm sampled across 4 separate sessions. In line with previous literature, we found that spots associated with cold sensations (112 cold spots, 34%) were more frequent than spots associated with warm sensations (41 warm spots, 12%). Still more frequent (165 spots, 49%) were spots that elicited inconsistent sensations when repeatedly stimulated by the same temperature. Remarkably, only 13 spots (4%) conserved their position between sessions. Overall, we show unexpected inconsistency of both the perceptual responses elicited by spot stimulation and of spot locations across time. These observations call to revise the traditional view that thermosensitive spots reflect the location of individual thermosensitive, unimodal primary afferents serving as specific labelled lines for corresponding sensory qualities.

**New & Noteworthy:** Thermosensitive spots are clustered rather than randomly distributed, and have highest density near the wrist. Surprisingly, we found that thermosensitive spots elicit inconsistent sensory qualities and are unstable over time. Our results question the widely believed notion that thermosensitive spots reflect the location of individual thermoreceptive, unimodal primary afferents, that serve as labelled lines for corresponding sensory qualities.

## Introduction

Thermoreception is not uniform across the skin surface.^1–5^ Even within a body part, there are small areas of unusually high thermal sensitivity, commonly referred to as ‘thermosensitive spots’.^6–23^ Early work reported that many spots were temperature-specific, eliciting either warm or cool sensations with the corresponding stimulus.^6^ Crucially, each spot was thought to indicate the presence of nerve endings from a single cutaneous afferent fibre, responding consistently to either warmth or cold.^17–23^ Thus, thermosensitive spots have provided foundational support for theories of neural specificity – the view that specific sensory qualities are associated with specific classes of afferent fibre.^24^ Later studies of the loss of sensation during pressure block and anaesthetic block showed that cold sensations were carried by thinly myelinated A -fibres, while warm sensations were carried by unmyelinated C-fibres, confirming the link between afferent fibre types and sensory qualities.^25^

Green and colleagues^11^ developed a two-step search method to identify thermosensitive spots across larger skin areas. Briefly, they used a thermode with a contact area of 16 mm^2^ to first identify broad thermosensitive sites, followed by a thermode with a contact area of 0.79 mm^2^ to identify the smaller, classical spots within those sites. They applied this procedure in the human forearm, classifying sites and spots according to the quality of the evoked sensations. They found that the quality of sensation evoked by a thermal stimulus could be inconsistent. Although 96.7% of sites remained sensitive over the experimental session, a surprising 31.8% were associated with different sensations across repeated tests, which presumably meant that their stimulations activated multiple thermosensitive primary afferents. In that case, smaller stimulation areas should produce more consistent sensory qualities – although this prediction was not tested in that study.

Such a study is required for two reasons. First, if thermosensitive spots are shown to be inconsistent and unstable over time, this might question the notion that each spot corresponds to a single afferent unit, since the skin locations of afferents’ nerve endings can be assumed to be unchanging. Second, near-threshold stimulation of a single thermosensitive spot can be considered to cause a minimal afferent signal to the brain. Neural specificity theories predict that even minimal afferent signals should consistently evoke the same sensation, because the “line” carrying the signal bears a “label” that is read by the brain as defining the sensory quality.

## Methods

### Subject details

8 participants (5 females; 18-35 years) were recruited from an institutional participant pool and compensated for their time. The sample size was chosen based on previous studies mapping suprathreshold thermosensitivity in the forearm.^3,16,26,27^ Participants with skin conditions or sensitivity skin were excluded. The experiment was approved by the UCL Research Ethics Committee.

Participants gave written consent to video recording and photography of their arm during the experimental session. They were invited to review recordings and images after the experiment.

### Experimental schedule

Our procedure to identify spots was based on the protocol described by Green et al.,^11^ but included several extensions and modifications. The procedure was repeated 4 times on different days. Sessions 1 and 2 were separated by 24 hours. In these 2 sessions, thermosensitive spots were identified based on detection of a warming stimulus 2°C above individual baseline skin temperature, or detection of a cooling stimulus 2°C below baseline. Sessions 3 and 4 took place 30 days after sessions 1 and 2 respectively, and used ±4°C variations. We predicted that larger temperature changes should reveal more thermosensitive sites, so this factor acted as an internal validation that our methods correctly tracked human thermosensitivity.

In each session, we used a two-step systematic search and classification procedure to identify thermosensitive spots (Figure 1). In Phase 1, we used a circular Peltier thermode (Physitemp NTE2A, diameter: 12.7 mm, contact area: 126.68 mm^2^) to search efficiently for general sites of high thermal sensitivity in the dorsal forearm. In Phase 2, we used blunted aluminium wires (diameter: 1 mm, contact area: 0.79 mm^2^) to scan for smaller thermosensitive spots within these larger sites (Figure 1). The data of interest here are the spots, with sites being just an intermediate step for efficient identification of spots. The blunted aluminium wires were maintained in a water bath (Premiere XH-1003, C&A Scientific Company, Virginia, USA Premiere) at the desired temperature. The experimenter held one end of the wire via a custom-made thermoinsulating handle.

**Figure 1.**
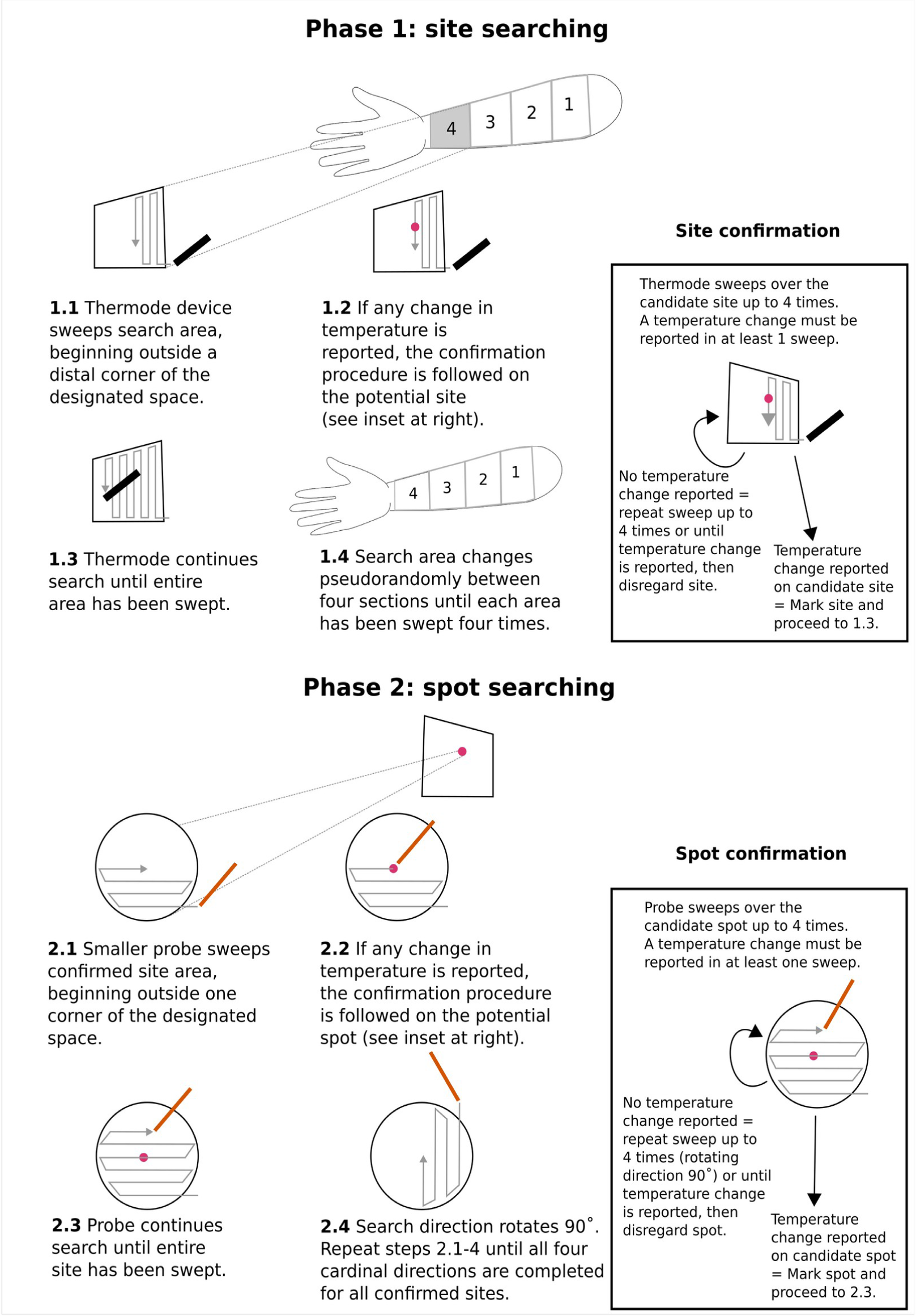
Spot searching method. In Phase 1, the dorsal forearm is divided into four equal segment and thermodes sweep each area to locate candidate thermosensitive sites. In Phase 2, each confirmed site is swept with an aluminium wire (contact area: 0.79 mm^2^) to locate thermosensitive spots.

The blunted aluminium wires did not have a closed-loop temperature control mechanism during spot search (Figure 1). Therefore, the temperature of the probe drifted towards room temperature once they were removed from the water bath. We calibrated this temperature drift using thermal imaging. To do so, we first measured the actual temperature of the wire probe after it had been warmed/cooled in a water bath by ± 4°C from a typical skin baseline value of 31°C. We found that the starting temperature of the wire was highly repeatable across two calibration sessions (calibration 1 (8 repetitions)-Cold mean: 26.8°C ± 0.09; Warm mean: 35.0°C ± 0.08 // calibration 2 (5 repetitions)-Cold mean: 27.0 ± 0.06°C; Warm mean: 35.1 ± 0.2°C).

Next, we measured how the thermal drift of the wire when it was swept across the skin to search for spots. From the start to the end of a sweep, cold wires changed by -0.44 ± 0.14°C (5 repeated sweeps), while warm wires changed by -1.80 ± 0.73 °C (5 repeated sweeps). The thermal energy of the warm stimuli is farther from room temperature, explaining the greater thermal drift. Crucially, the thermal drift did not reach or cross the baseline temperature of the skin for neither the warm nor the cold stimuli. Thus, effective thermal stimulation was present throughout the sweep.

Laboratory room temperature was maintained at 23°C by an air conditioning unit. The experiment was recorded with a 720×720 pixel camera located 53 cm above the table, giving an effective spatial resolution of 0.33 mm/pixel. The table was covered with 1-mm graph paper allowing accurate repositioning of the arm, and thus comparison of spot locations across sessions.

### Procedure

After obtaining informed consent, the right forearm was placed comfortably on the table, with the dorsal side upwards. To familiarise participants with the sensations they should report, we demonstrated and narrated the procedure for locating a single site (Phase 1). Participants were instructed to report immediately by saying “warm” or “cold” if they felt any change in the temperature of the applied thermal probe.

Participants were then blindfolded. The tip of the middle finger and centre of the elbow were aligned to the graph paper. The distance from the wrist to elbow was measured and the forearm divided into four equal segments, which were marked on the paper and visible to the camera. The graph paper from the first session was kept for each individual to allow precise repositioning in future sessions, and standardisation of coordinates for image alignment and analysis.

Thermal stimuli were specified relative to each participant’s baseline skin temperature at the beginning of each session. Using a laser thermometer, skin temperature was measured adjacent to the wrist and elbow. The cooling stimulus was set to either 2°C (sessions 1,2) or 4°C (sessions 3,4) below the lower of the these and warming stimulus was set to 2/4°C above the higher of the same two temperatures. Cold and warm stimuli were tested in separate, counterbalanced blocks within each session.

In Phase 1, the four areas of the forearm were tested in pseudorandomised order to prevent both order effects and temporal summation.^28,29^ In each area, thermosensitive sites were located by sliding the thermode over the skin. A silicone-based lubricating gel was applied to minimise friction and excessive mechanorecptor stimulation during movement of thermode. The weight of the thermode provided the downward force: the experimenter exerted no additional pressure. The thermode was placed in one corner of each area and systematically swept across it in a medio-lateral direction (Figure 1). Each area was searched four times. At the end of each medio-lateral sweep, the thermode was moved proximally to begin the next sweep. The sweeps began and ended just outside the boundaries of each of the four area to prevent onset/offset effects (Figure 1).

If participants reported “warm” or “cold” sensations at any point during a search, this was considered a candidate thermosensitive site. We marked the location on the skin with coloured ink, and followed by sweeping up to four further times to confirm the site (Figure 1). These follow-up sweeps could help distinguish genuine thermal sensations from potential false-positive reports. If participants reported any thermal sensation during any follow-up sweep, then the location was marked as confirmed thermosensitive site, and the confirmation procedure was terminated. Importantly, the reported sensations did not need to be consistent with the actual stimulus temperature, nor with each other. If no thermal percept was reported in any of four confirmation sweeps, the candidate site was classed as unconfirmed.

In Phase 2, we then searched for smaller thermosensitive spots within each confirmed site, by repeating at a smaller scale the same process used to search for sites. This time we rotated the direction of each successive confirmation sweep by 90 degrees in order to discourage participants from responding simply on the basis of memory for elapsed time or for tactile location. In place of thermodes, we now used much smaller warmed or cooled aluminium wire as stimulators (Figure 1).

At the beginning of a search, the experimenter took one of the aluminium wires in the thermal bath from the custom-made thermoinsulating handle. Then, the experimenter dried excess water with absorbent tissue and began to search for spots within the larger site. Contact with the skin was made within about 2 s of the removal of the wire from the water bath. The sweep lasted until a spot was reported or until the entire site was swept, which took approximately 7 s (16 mm^2^). After every sweep or spot location, the experimenter placed the probe back into the water bath. We had multiple identical probes in the water bath. The experimenter alternated between the probes to allow each probe to return to the bath temperature before being used again.

When a spot was located and subsequently confirmed (Figure 1), it was marked on the skin. If a participant consistently reported a temperature sensation corresponding to the stimulus temperature (i.e., ‘cold’ to temperature 2/4°C below baseline and ‘warm’ to temperature 2/4°C above baseline) both on initial identification and subsequent confirmation, then the spot was classified as cold or warm. If a participant reported different temperature sensations when the potential spot was first identified and in any of up to four confirmation attempts, then the spot was classified as inconsistent. Spots that elicited sensations to both stimulus temperatures in separate blocks were classified as inconsistent. Occasionally, initial identification and subsequent confirmation responses were consistent with each other, but did not correspond to the actual stimulus temperature: these spots were classified as incongruous (Figure 2A). Warm, cold, inconsistent and incongruous spots were marked on the skin with four different ink colours. Some spots initially yielded a thermal sensation, but no further sensation was reported on any of four subsequent stimulation confirmation attempts with the same stimulus. These spots were considered unconfirmed and were identified with a different ink. At the end of each session, a final image was taken of the positions of all spots.

**Figure 2.**
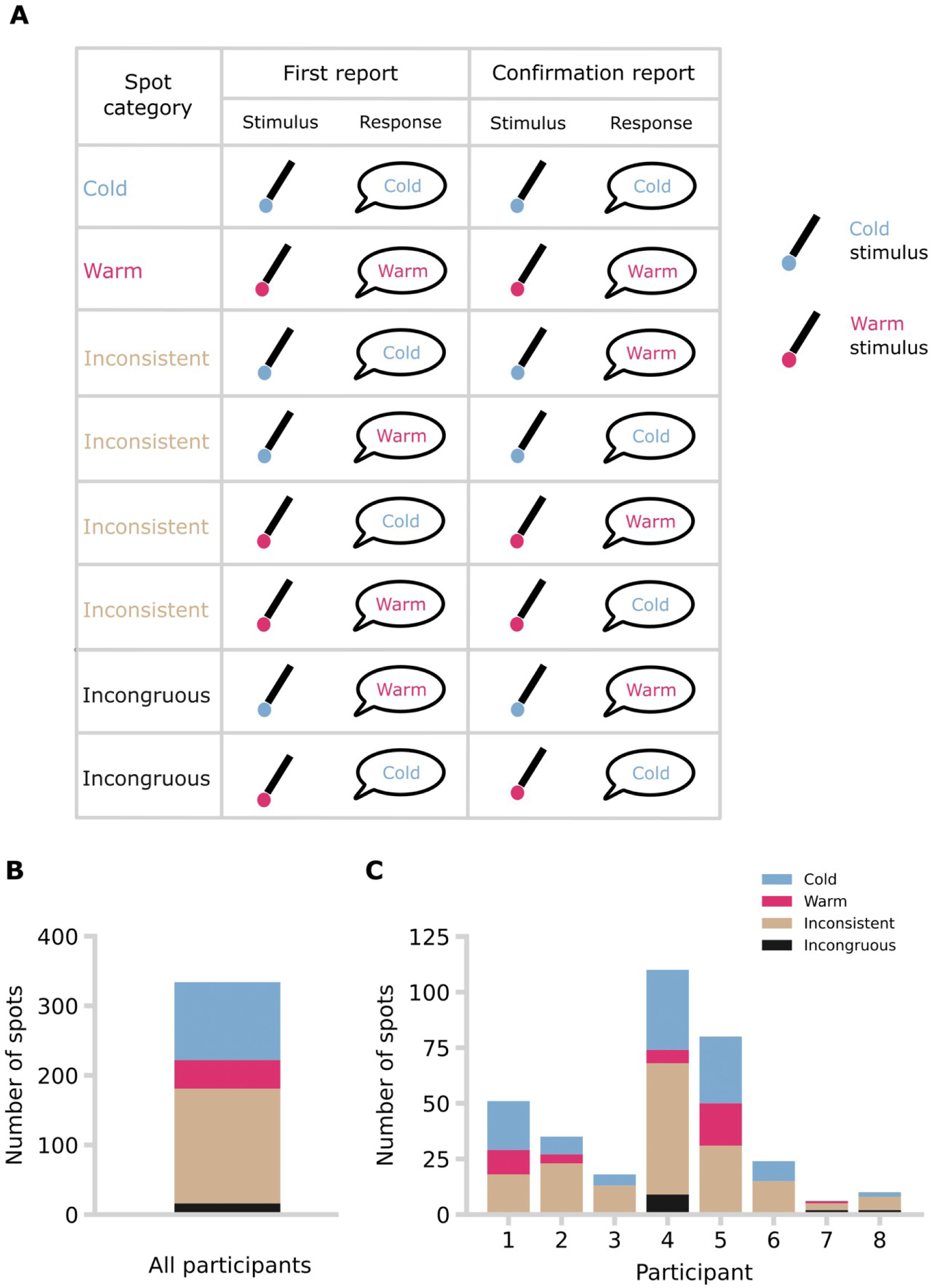
Classification and distribution of spots by sensation elicited, with respect to modality of stimulus. **A)** A table with the taxonomy of spots is shown. **B)** Total number of spots across participants by spot category. **C)** Total number of spots per participant and by spot category.

### Analysis

The final images of each session were pre-processed. First, skin markings were annotated with a graphics editing program. Second, the images within each participant were aligned across sessions with DS4H Image Alignment^30^ by defining a few fiducial points. Third, spot location data was extracted from these standardised images with a custom Python script (see software repository: https://github.com/iezqrom/publication-thermal-spots-quality-location-inconsistent).

Briefly, the centre of the digital mark assigned to each spot was manually clicked and an XY coordinate recorded. Forearm curvature was ignored. The classification of each spot was saved with the coordinates.

Spot classifications were compared across sessions and subjects. For some analyses, parametric or non-parametric tests were chosen depending on data normality. Unconfirmed spots were not included in this and subsequent analysis.

To assess spatial distribution of spots along the forearm, we used the Anderson-Darling test^31^ to test for a uniform distribution of the spots’ X-coordinates between elbow and wrist. The uniform distribution tested had a lower bound of 0 and an upper bound of 1200 pixels. We focussed on this spatial axis because thermosensitivity shows a proximo-distal gradient,^3,5^ and because this axis was less affected by curvature distortions that would affect mediolateral position estimates. Data from each participant was tested separately, but data were pooled across sessions. Deviation from a uniform distribution would indicate that spots are more likely to be reported in certain locations on the dorsal forearm (for example, near the wrist, or elbow). Spot data were pooled across all four sessions. One participant reported only six spots, which was insufficient to estimate distribution, and was thus excluded from this test.

We also quantified spatial aggregation of spots. We compared the distance from each spot to its ‘nearest neighbour’ using the Clark-Evans Aggregation Index, *R*.^32^ As there could be additional spots outside of our measured boundaries^13^, we applied a correction for edge effects.^33^ Spot data were pooled across all sessions.

To estimate stability and consistency of thermosensitive spots, we next compared the spatial positions of spots in each session with those in all other sessions within each participant. Repeatable repositioning of the arm is clearly crucial for this analysis, and we applied several strategies to standardise forearm positioning (see Procedure). Additionally, we performed image alignment. A spot was considered conserved if any spot in any other session was less than 2 mm (6 pixels) away. This criterion was based on twice the diameter of the aluminium wire used for stimulation.

## Results and discussion

### The sensory quality evoked by spot stimulation is variable

We extended Green’s method^11^ for studying thermosensitive spots (Figure 1), using repeated systematic searches over a large skin region (the entire forearm), at extended timescales (days and months). We identified a total of 349 spots across participants of which 334 (mean = 10.44 ± 10.63 SD) were confirmed following the confirmation procedure (Figure 2A). Only confirmed spots were included in subsequent analyses. Crucially, we then distinguished between spots that consistently elicited a single sensory quality of warmth or cold on repeat testing, and inconsistent spots that evoked different sensory qualities when repeatedly tested with the same thermal stimulus.

Consistent with previous work,^6–8,10,11^ spots eliciting ‘cold’ responses (n = 112, mean = 14.00 ± 13.55 SD) were more frequent than those eliciting ‘warm’ responses (n = 41, mean = 5.13 ± 6.81 SD W = 35.00, *p* < 0.01, r = 0.944, Wilcoxon signed-ranks test). We found 165 inconsistent spots, which amounts to 49% of all confirmed spots. Thus, the inconsistency of evoked sensory qualities reported by Green and colleagues^11^ for much larger thermal sites of 16 mm^2^ was found also for much smaller thermosensitive spots of just 0.79 mm^2^. Crucially, we found more spots when we used more extreme temperatures (±2°C-total spots: 148, mean = 18.5 ± 18.3; ±4°C-total spots: 186, mean = 23.25 ± 19.1), suggesting our thermal stimulation was functional and working as expected.

### Spots are aggregated and non-uniformly distributed

Thermosensitive spots have classically been taken as a proxy of the anatomical distribution of thermosensitive afferent innervation. However, studies of spot spatial distribution have been limited to small subregions of the hand or forearm ^6–18^. Green et al. (2008)^11^ searched for spots across the entire forearm, but did not analyse their spatial distribution properties. This data would contribute to our understanding of the relationship between spots and thermosensitive afferent innervation.

Visual inspection of our data shows that spots were distributed unevenly across the forearm (Figure 3A). We applied three different analyses to describe the spatial properties of spots. First, the distribution of spots deviated significantly from a uniform spatial distribution for four out of the seven participants included in this analysis (Figure 3A). Second, dividing the forearm into four equal distal-proximal areas showed no significant main effect, nor interaction effect, in spot density (F _3, 28_ = 2.14, p = .118, ɳ_p_^2^ = 0.19) (Figure 3B), ruling out a simple spatial gradient hypothesis, though visual inspection shows a relatively high density of spots close to the wrist. Third, the Clark-Evans Aggregation Index was significantly below 1 for all participants tested, providing strong evidence of spot aggregation (Figure 3C). Altogether, these results show that the spatial distribution of spots was non-uniform and followed an aggregated pattern. Additionally, spots were most frequent just proximal to the wrist, but did not follow any obvious proximodistal gradient.

**Figure 3.**
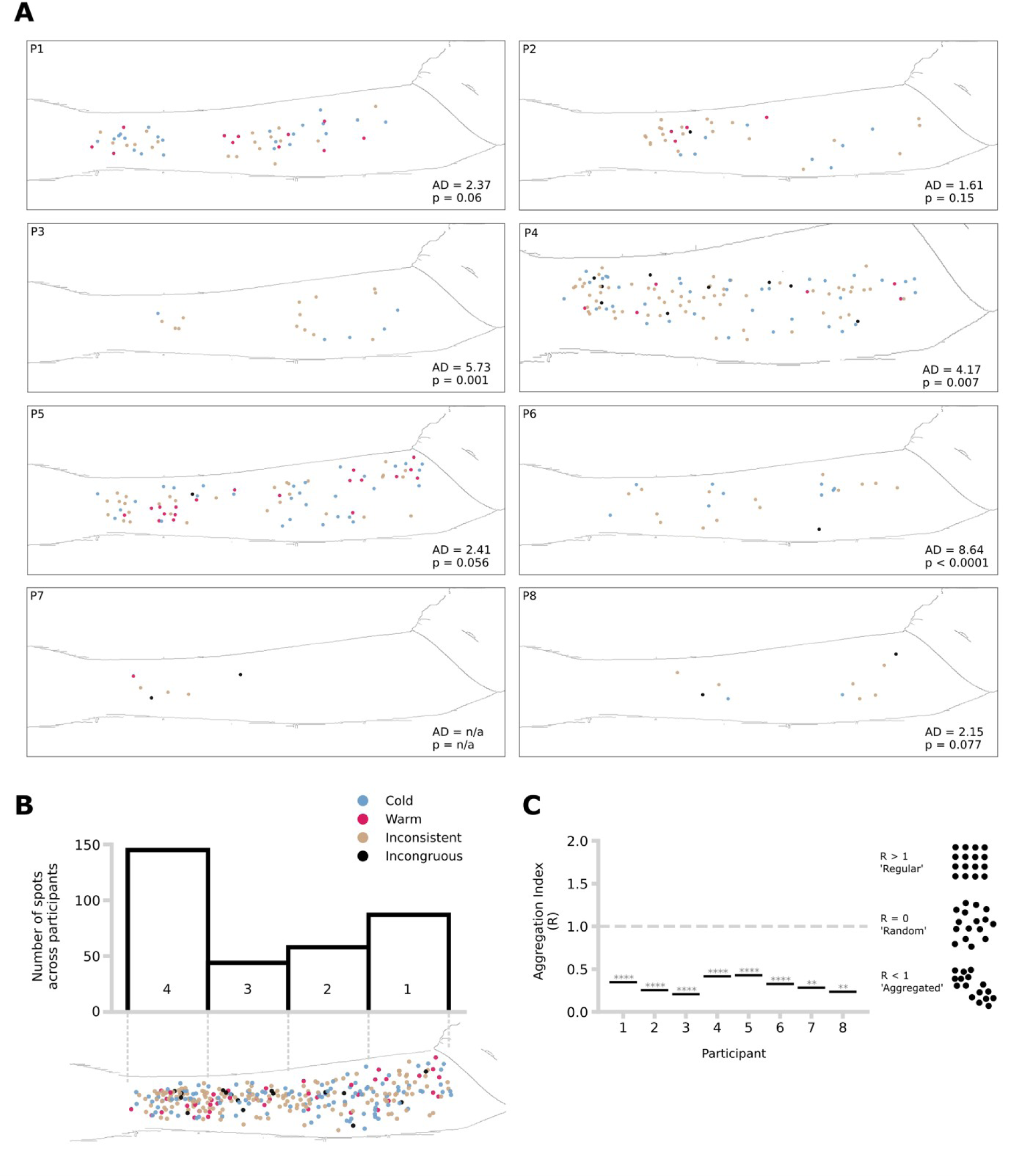
Spot spatial distribution. **A)** Spot distribution across participants. A single forearm silhouette has been placed in each box for visualisation purposes only. Anderson-Darling (AD) test results and associated p-values are shown in each panel at the bottom right corner. **B)** Total number of spots pooled across participants by search area. The top panel shows the number of <u>spots per skin search area (1-4) across all participants and sessions. The bottom panel is a</u> visualisation of the distribution of all spots across participants and sessions in a template forearm silhouette. **C)** Aggregation index (Clark-Evans aggregation index, R) of confirmed spots per participant, with Donnelly correction. Illustrative examples are shown on the right. Asterisks indicate the p-values obtained from two-sided test statistics. ** p < .01, **** p < .0001.

### The location of spots varies across testing sessions

If spots reflect the presence of nerve endings that are stable, then the same spots should be found across repeated searches.^8,12^ However, no study has addressed this question with repeated systematic searches over large skin regions.

We found that conservation of spots across testing sessions was very rare (Figure 4). Just 13 of 334 confirmed spots were re-identified between sessions. Of the 13 conserved spots, 11 had the same classification (inconsistent/warm/cold) across <u>sessions. No spot was conserved across 3 or more sessions.</u>

**Figure 4.**
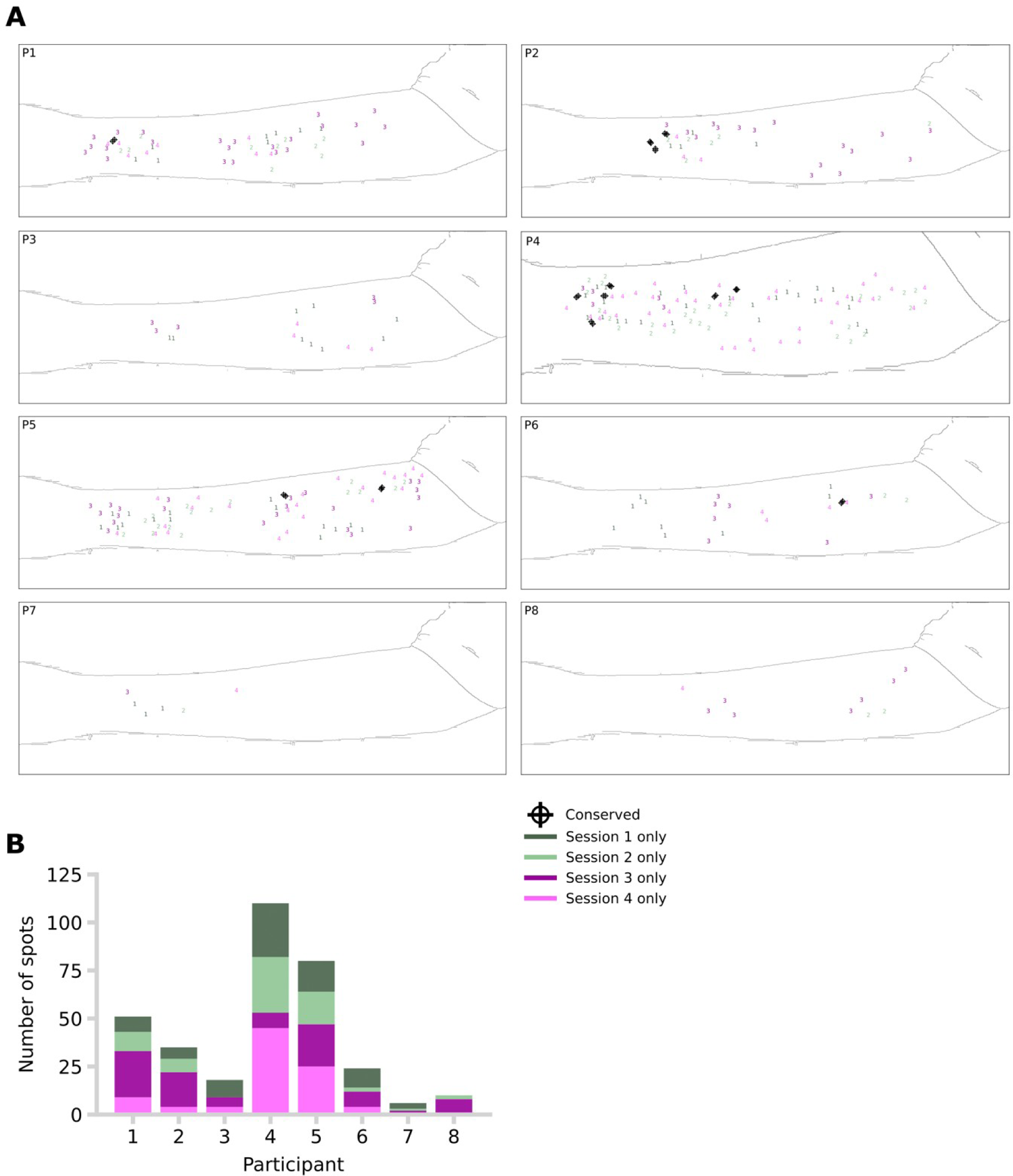
Conservation of spots. **A)** Position of spots per participant and session. The spots that were considered conserved across sessions are indicated with a black dot and cross. A single forearm silhouette has been placed in each box for visualisation purposes only. **B)** Total number of spots per participant and session.

## Discussion

We investigated the quality and spatiotemporal features of thermosensitive spots on the human forearm, extending previous studies^11,6,7,14^. We confirmed the presence of 334 thermosensitive spots across 8 participants. We found more cooling-than warming-responsive spots across all participants. Surprisingly, we found 165 spots (49%) of spots elicited inconsistent reports of perceived thermal quality. That is, repeated identical temperature stimulation of the same spot would produce both ‘cold’ and ‘warm’ responses. The spatial distribution of the spots was non-uniform and followed an aggregated pattern. Spots were most frequent just proximal to the wrist, but did not follow any obvious proximodistal gradient. Finally, we observed a surprisingly low conservation rate over time: only 4% were reidentifiable on successive sessions.

We found more cold-sensitive spots (34%, n = 112) than warm-sensitive spots (12%). Previous studies have also found more spots eliciting ‘cold’ than a ‘warm’ responses^6–8,10,11^, but we cannot directly compare the type and frequency of spots because of differences in body region, stimulus size, thermal magnitude, and search protocol. Based on our data and previous studies, we also cannot conclude that there are more cold-sensitive than warm-sensitive spots for three reasons. First, humans are more sensitive to cooling than to warming. In other words, the relative temperature change required to detect a cooling stimulus is smaller than the temperature change required to detect a warming stimulus^1^. Second, the endings of cold-sensitive fibres are found more superficially than the endings of warm-sensitive fibres.^34–36^ Third, some cold-sensitive fibres are Aδ-fibres, whereas all warm-sensitive fibres are C-fibres with slower conduction velocities^37–40^. The combination of these factors may mean that less warm-sensitive spots were detected in our study and others because processing warm signals takes longer and is noisier than processing cold signals. In our study, we used the same magnitudes (±2°C & ±4°C) for cold- and warm-sensitive spot search, which may have biased the frequency of spot type against warm-sensitive spots. Future studies could address the question whether there are more cold-than warm-sensitive spots by matching the magnitude of the thermal stimuli to account for differences between cold- and warm-sensitive neural circuits.

The number of spots that elicited inconsistent reports of perceived thermal quality was high. This seems at odds with the way that thermosensitive spots have classically been interpreted. In particular, our results question the repeated notion that thermosensitive spots reflect the location of individual thermoreceptive primary afferents,^16–23^ that serve as labelled lines for corresponding sensory qualities. Our stimulator (contact area: 0.79-mm^2^) might have stimulated a multimodal primary afferent, rather than a non-noxious, unimodal thermoceptive afferent. Since polymodal fibres, by definition, are activated by multiple stimulus types and do not carry a distinctive stimulus quality, their recruitment could potentially explain our inconsistent responses. There are two types of multimodal afferents to consider in <u>our study.</u>

First, tactile signals might prime or modify thermal signals. We minimised multimodal, thermotactile stimulation by reducing friction with lubricant, but there would still be some tactile pressure signals encoded by slowly-adapting (SA1, SA2) and intermediate-adapting (C-tactile) afferents in the skin. These afferent types have <u>been shown to change firing with sustained pressure and thermal changes,</u> potentially contributing to thermal sensations in unknown ways^41,42^. Second, warm and cold sensations might be mediated by multimodal C-fibres. Traditionally, innocuous cold sensations are thought to be mediated by Aδ-fibres, while innocuous warm sensations are mediated by C-fibres.^34,37,38^ The responses of these fibres are driven by TRPM8 receptor channels in cooling-responsive afferents and by TRPV1 in warming-responsive fibres on warming.^34,38^ However, a microneurography study showed that cold-sensitive C-fibres responded both to cold and warm stimuli.^43^ Consistent with this finding, mice without the cooling-sensitive receptor, TRPM8, are unable to perceive warm.^39^ Thus, a specific sensory quality may depend on polymodal afferents, rather than specific afferents, contrary to labelled-line theories.^24^ Interestingly, recent models of tactile afferent coding^44,45^ have also relinquished the strong assumption of labelled-line coding that underlay classical models.^46^ If sensory quality is mediated by polymodal afferents, this could be a source of variability in evoked sensations, particularly when a single afferent is stimulated.

Intraneural microstimulation potentially provides direct tests of the relation between specific afferents and a sensory quality. Such stimulation bypasses the transduction process at the peripheral receptor, by stimulating the afferent directly. Microneurography studies have shown that stimulation of single primary afferents reliably produces a localised, distinct and pure sensory quality, though this conclusion is based on mechanosensitive Aβ-fibres rather than thermosensitive Aδ- or C-afferents.^47^ Nevertheless, if we assume that our stimuli activated a single thermosensitive fibre, then we can suggest either that the inconsistent sensory <u>qualities observed in our study might arise in the process of transduction at the</u> receptors, or that the concept of an individual labelled line for sensory quality is incorrect.

Our current design focusses on minimal sensations with small, near-threshold stimuli. Classically, these sensations were attributed to a single primary afferent. However, we do not have neurophysiological evidence to confirm this assumption. We can be confident that we indeed stimulated thermal afferents, because we found more spots in testing sessions using more extreme thermal stimuli. However, during searching for spots, we may have stimulated receptive fields of two or more afferents that overlap in the same skin location. While we cannot rule out this possibility, it still seems surprising that the sensory quality evoked by repeated stimulations was so often inconsistent. The challenge from spot inconsistency to the concept of labelled <u>lines remains.</u>

Alternatively, the frequent inconsistency we found could reflect a central sensory process that misreads unreliable thermal input from one or more primary afferents. For instance, a recent study found that larger thermal stimuli produce psychophysical functions with higher precision than smaller stimuli, suggesting that averaging over multiple afferents reduces sensory noise.^48^ Population coding, in which sensory quality depends on a balance of activity across many different afferents, potentially differing in physiological type as well as in location, may play a crucial role in robust and stable thermosensation.^49^ In the thermal system, spatial summation is a well-known feature in both object-level perception and in thermoregulation.^50,51^ In our <u>study, we use small probes to study thermosensation in its role during object-level</u> perception. However, we do not know the minimal primary afferent activity required to detect a thermal sensation.

A seminal study by Johnson & Darian-Smith^52^ about warmth intensity discrimination suggested that, for warmth discrimination, the combined input of ∼20 fibres is required to match human performance with cortical responses in monkeys. Crucially, this conclusion is based on correlating monkey neuron recruitment data with human performance. This study is effectively about suprathreshold intensity coding, as might be tested in psychophysical scaling studies. It does not state that ∼20 fibres are necessary to have a thermal sensation, but that ∼20 fibres are sufficient to reconstruct the range of thermal intensity perception.^52^ Interestingly, a recent study of visual sensory qualities reported that simulation of a single retinal M-cone in vivo could often produce an achromatic percept^53^ – a striking finding given that colour vision has been the paradigmatic evidence for labelled lines. This study, like ours, suggests that a minimal afferent signal may be insufficient to evoke a sensory quality. Presumably some element of evidence accumulation across time or across multiple afferent fibres is required for a stable sensory quality – a quantum for qualia. In that case, Muller’s original metaphor of a *label*, i.e., a self-intimating sensory quality based on the origin of each neural signal, should be discarded.

Consistent with previous research on the insensitivity to warmth in subregions of the forearm,^10^ we found that spots tended to aggregate across the forearm (Figure 3). We also report significant non-uniformity in spatial distribution, with more spots observed closer to the wrist (Figure 3). Our results are seemingly inconsistent with previous mapping studies. Specifically, we found a higher number of spots distally within the forearm whereas previous studies have shown a proximodistal decrease in thermal and pain sensitivity^1,3,4,54^. However, these previous studies have compared thermal sensitivity across the entire body. The proximodistal gradient that they report was based on contrasting the torso and the extremities. Importantly, our high-density thermosensory data shows there is a relative increase in thermal sensitivity around the wrist area^3,4^. Our study is thus compatible with previous perceptual studies of other sensory modalities, and shows for the first time the spatial distribution of spots following a systematic search across a large skin region. Future studies should systematically search for spots across the entire body and compare distribution across body sites.

We found a low conservation rate of spots (4%) across days and weeks. We advance three possible alternative explanations for the surprising instability. First, sensory detection reports may depend heavily on context, including experience prior to each session. Context-dependent sensitivity is known to be important in sensations at noxious temperatures,^55,56^ but may also apply also to the non-noxious temperatures studied here. Second, fluctuations of peripheral excitability across time may also play a major role in thermoception.^57^ For instance, thermal detection thresholds have been found to vary by 0.9°C in the hand of healthy young adults. Third, tactile afferent innervation renews throughout an animal’s lifetime,^58^ but the rate of renewal of thermosensitive innervation in humans is unknown. Our observations were necessarily limited to the roughly 90 minutes of individual sessions, and the 31 days that separated the first from the last session. However, we found minimal conservation of spots even between sessions separated by just 24 <u>hours. Wholesale changes in the presence and location of receptor structures over</u> such short timescales seem unlikely. Therefore, we suggest that non-conservation reflects some process as yet unknown. Future studies should map thermosensitive spots over a wider range of time intervals, with a particular focus on repeat testing at regular intervals up to 1 day. A more comprehensive sensitivity profile might reveal a clearer picture of time-varying sensitivity. Optical Coherence Tomography^59^ promises the possibility of longitudinal imaging of sensory afferent fibres in vivo in future studies.

The low conservation rate could reflect methodological limitations when aligning the arm or spatial data. If our low conservation were due to these technical issues, visual inspection would show a common spatial pattern of spots within each session, which is simply shifted between sessions due to misalignment. We saw no evidence for this (Figure 4A). Similarly, mere misalignment would imply equal numbers of spots in each session. However, the number of spots varied across sessions as well as their locations (Figure 4B). The low conservation of spots across sessions is therefore unlikely to be due to limitations in arm positioning or data alignment.

A poor signal to noise ratio in thermal afferents would also lead to low measures of conservation. A spot might be identified on one session, but missed on another simply because of fluctuations in combined signal and noise reaching a central site for decision-making. However, high noise levels would imply a high false negative rate with stimulations of an afferent fibre often producing no thermal sensation (SDT misses). In our dataset, unconfirmed spots can be taken as a proxy for such false negatives. However, only 15 spots out of a total of 349 (4.3%) identified were classified as unconfirmed, a value similar to previous research.^11^ Therefore, it is unlikely that methodological issues or sensory noise can account for low rates of conservation.

Our stimulator for spot search was not temperature-controlled, and maintaining temperature stability of probes during dynamic skin contacts is challenging. ^60^ Therefore, the high rate of inconsistency could be due to low repeatability and stability of the thermal stimulus used for spot search. We think this is unlikely for three reasons. First, we used a temperature-controlled probe for our initial search for larger thermosensitive sites, and we only searched for spots within such confirmed sites. Second, we found more spots when we used more extreme temperatures, which is expected as greater stimulus amplitudes are more likely to reach detection thresholds. Third, we showed that the starting temperature of our small stimulator was consistent. Importantly, we showed that the thermal shift during the stimulation period itself was repeatable, and could not explain the inconsistency in the quality of the evoked sensations. This makes it unlikely that our finding of frequent inconsistent spots merely reflects ineffective stimulation. Interestingly, Green and colleagues^11^ also reported inconsistency of evoked sensory qualities with large, temperature-controlled thermodes (contact area: 16 mm^2^). In our study, we report inconsistency of the evoked sensory qualities, and, for the first time, instability of spatial location of thermosensitive spots.

Both the inconsistency of sensory qualities and the spatial instability of spots are likely to have a neurophysiological or perceptual origin. A limitation of our protocol is that we used the same stimulus temperature for the entire forearm. We adjusted the temperature of the thermal stimulus to each participant’s baseline temperature after a period of acclimatization by measuring the temperature of two points in the skin. However, skin temperature is not homogenous across the skin.^61^ Future studies should combine online thermal measurements with feedback control to adjust stimulus temperature according to the baseline temperature of the stimulated skin region and better describe the relationship between the magnitude of thermal stimulation and the number of identified spots.

In our study, we observed a surprising interindividual variability in the number of confirmed spots. Previous studies have reported substantial interpersonal variability in thermosensitivity,^3,4^ but individual differences in thermosensitive spot distribution have not been studies systematically, to our knowledge. The interpersonal variability we observed could be due to different factors such as genetic, hormonal or perceptual characteristics. Our study was not designed for investigating individual differences, but focussed on obtaining systematic and common patterns in the spatiotemporal characteristics of spots. Moreover, our dataset is limited for making conclusions about the absolute numbers of spots in the human skin. First, although the sample size in our study is similar to previous studies on suprathreshold thermosensitivity in the forearm^3,16,26,27^, the number of spots and participants in our dataset is not sufficient to make strong claims about individual differences and about the frequency of spots at a population level. Additionally, we only studied one body site-the forearm. Thermal sensitivity varies across body regions^1,3,4^. Therefore, the distribution of spots may differ between body sites. The design of our study was suitable for finding differences in the distribution of spots spatially and temporally within a body site. Future studies should characterise the types and frequencies of spots over a larger sample with different populations and across multiple body regions.

Overall, our study confirms the existence of thermosensitive spots, consistent with previous studies.^6,7,11^ However, we found that these spots often produced inconsistent sensory qualities, and were unstable over time. Our results call into question the widespread notion that thermal spots indicate the presence of individual thermosensitive primary afferents projecting centrally as labelled lines, and that minimal activation of an individual labelled line is sufficient for the distinct and reliable phenomenal experience of a specific sensory quality. Our results do not rule out some form of neural specificity theory at the level of fibre populations, but they do suggest that label metaphors for sensory quality should be revised.

## Supplemental materials

Raw data and source code can be found in the following repository: iezqrom/publication-thermal-spots-quality-location-inconsistent: Code & data supporting academic publication “The sensory quality and spatiotemporal location of thermal spots are inconsistent.” published at TBD (github.com)

## Acknowledgements

We would like to thank Dr. Caterina Maria Leone and Dr. Antonio Cataldo for the discussions at the inception of the project. We are grateful for the advice of Prof. John Mollon.

## Author contributions

Conceptualization: P.H.; Methodology: I.E.R, M.F.C. and P.H.; Software: I.E.R; Validation: I.E.R and P.H.; Formal Analysis: I.E.R and M.F.C.; Investigation: I.E.R, M.F.C., S.C. and P.H.; Resources: P.H. and G.D.I.; Data Curation: I.E.R, M.F.C. and S.C.; Writing – Original Draft: I.E.R and P.H.; Writing – Review & Editing: I.E.R and P.H.; Visualization: I.E.R and M.F.C.; Supervision: S.C., G.D.I and P.H.; Project Administration: S.C. and P.H.; Funding Acquisition: G.D.I and P.H.

## References

1. Stevens, J. C., & Choo, K. K. (1998). Temperature sensitivity of the body surface over the life span. Somatosensory & Motor Research, 15(1), 13–28. 10.1080/08990229870925

2. Agostino, R., Cruccu, G., Iannetti, G., Romaniello, A., Truini, A., & Manfredi, M. (2000). Topographical distribution of pinprick and warmth thresholds to CO2 laser stimulation on the human skin. Neuroscience letters, 285(2), 115–118. 10.1016/s0304-3940(00)01038-7

3. Filingeri D., Zhang H., & Arens E. A. (2018). Thermosensory micromapping of warm and cold sensitivity across glabrous and hairy skin of male and female hands and feet. Journal of Applied Physiology, 125(3), 723–736. 10.1152/japplphysiol.00158.2018

4. Luo, Maohui, Zhe Wang, Hui Zhang, Edward Arens, Davide Filingeri, Ling Jin, Ali Ghahramani, Wenhua Chen, Yingdong He, and Binghui Si. (2020). High-density thermal sensitivity maps of the human body. Building and environment, 167, 106435. 10.1016/j.buildenv.2019.106435

5. Li, X., Petrini, L., Defrin, R., Madeleine, P., & Arendt-Nielsen, L. (2008). High resolution topographical mapping of warm and cold sensitivities. Clinical neurophysiology, 119(11), 2641–2646. 10.1016/j.clinph.2008.08.018

6. Blix, M. (1882). Experimentela bidrag till lösning af frågan om hudnervernas specifika energi. Uppsala Läkför Förh, 18, 87–102. 10.1016/s0361-9230(98)00067-7

7. Donaldson, H. H. (1885). On the temperature-sense. Mind, 10(39), 399–416. https://psycnet.apa.org/doi/10.1037/11304-035

8. Dallenbach, K. M. (1927). The temperature spots and end-organs. The American Journal of Psychology, 39(1/4), 402–427. 10.2307/1415426

9. Burnett, N. C., & Dallenbach, K. M. (1927). The Experience of Heat. The American Journal of Psychology, 38(3), 418–431. 10.2307/1415010

10. Green, B. G., & Cruz, A. (1998). ‘Warmth-insensitive fields’: Evidence of sparse and irregular innervation of human skin by the warmth sense. Somatosensory & Motor Research, 15(4), 269–275. 10.1080/08990229870682

11. Green, B. G., Roman, C., Schoen, K., & Collins, H. (2008). Nociceptive sensations evoked from ‘spots’ in the skin by mild cooling and heating. Pain, 135(1), 196–208. 10.1016/j.pain.2007.11.013

12. Norrsell, U., Finger, S., & Lajonchere, C. (1999). Cutaneous sensory spots and the “law of specific nerve energies”: History and development of ideas. Brain Research Bulletin, 48(5), 457–465. 10.1016/S0361-9230(98)00067-7

13. Strughold, H., & Porz, R. Die Dichte der Kaltpunkte auf der Haut des menschlichen Körpers. Zeitschrift für Biologie, 1931, 91,563–571.

14. Jenkins, W. L. (1939). Studies in thermal sensitivity: 9. The reliability of seriatim cold-mapping with untrained subjects. Journal of Experimental Psychology, 24(3), 278. https://psycnet.apa.org/doi/10.1037/h0056425

15. Jenkins, W. L. (1939). Studies in thermal sensitivity: 10. The reliability of seriatim warm-mapping with untrained subjects. Journal of Experimental Psychology, 24(4), 439. https://psycnet.apa.org/doi/10.1037/h0063226

16. Yang, F., Chen, G., Zhou, S., Han, D., Xu, J., & Xu, S. (2017). Mapping sensory spots for moderate temperatures on the back of hand. Sensors, 17(12), 2802. 10.3390/s17122802

17. Hensel, H., & Schafer, K. (1984). Thermoreception and temperature regulation in man. In Recent advances in medical thermology (pp. 51–64). Springer, Boston, MA.

18. Kenshalo, D. R., & Gallegos, E. S. (1967). Multiple temperature-sensitive spots innervated by single nerve fibers. Science, 158(3804), 1064–1065. 10.1126/science.158.3804.1064

19. Jones, L. A., & Ho, H. N. (2008). Warm or cool, large or small? The challenge of thermal displays. IEEE Transactions on Haptics, 1(1), 53–70. 10.1109/TOH.2008.2

20. Kenshalo, D. R., & Gallegos, E. S. (1967). Multiple temperature-sensitive spots innervated by single nerve fibers. Science, 158(3804), 1064–1065. 10.1126/science.158.3804.1064

21. Spray, D. C. (1986). Cutaneous temperature receptors. Annual review of physiology, 48(1), 625–638. 10.1146/annurev.ph.48.030186.003205

22. Hensel, H., & Iggo, A. (1971). Analysis of cutaneous warm and cold fibres in primates. Pflügers Archiv, 329(1), 1–8. 10.1007/BF00586896

23. Hensel, H., Andres, K., & Düring, M. V. (1974). Structure and function of cold receptors. Pflügers Archiv, 352(1), 1–10. 10.1007/BF01061945

24. Müller, J. P. (1833/1840). Handbuch der Physiologie des Menschen, 2 volumes, J. Hölscher, Coblenz. Trans. by William Baly (1838/1842) Elements of Physiology. In London: Taylor and Walton.

25. Mackenzie, R. A., Burke, D. A. V. I. D., Skuse, N. F., & Lethlean, A. K. (1975). Fibre function and perception during cutaneous nerve block. Journal of Neurology, Neurosurgery & Psychiatry, 38(9), 865–873. 10.1136%2Fjnnp.38.9.865

26. Molinari, H. H., Greenspan, J. D., & Kenshalo, D. R. (1977). The effects of rate of temperature change and adapting temperature on thermal sensitivity. Sensory processes, 1(4), 354–362.

27. Montgomery, L. D., & Williams, B. A. (1976). Effect of ambient temperature on the thermal profile of the human forearm, hand, and fingers. Annals of biomedical engineering, 4(3), 209–219. 10.1007/BF02584515

28. Stevens, J. C., Okulicz, W. C., & Marks, L. E. (1973). Temporal summation at the warmth threshold. Perception & Psychophysics, 14(2), 307–312. https://psycnet.apa.org/doi/10.3758/BF03212396

29. Banks, W. P. (1976). Areal and temporal summation in the thermal reaction time. Sensory processes, 1(1), 2–13.

30. Bulgarelli, J., Tazzari, M., Granato, A.M., Ridolfi, L., Maiocchi, S., De Rosa, F., Petrini, M., Pancisi, E., Gentili, G., Vergani, B., et al. (2019). Dendritic cell vaccination in metastatic melanoma turns “non-T cell inflamed” into “T-cell inflamed” tumors. Frontiers in immunology, 2353. 10.3389/fimmu.2019.02353

31. Anderson, T. W., & Darling, D. A. (1952). Asymptotic Theory of Certain ‘Goodness of Fit’ Criteria Based on Stochastic Processes. The Annals of Mathematical Statistics, 23(2), 193–212. 10.1214/aoms/1177729437

32. Clark, P. J., & Evans, F. C. (1954). Distance to Nearest Neighbor as a Measure of Spatial Relationships in Populations. Ecology, 35(4), 445–453. 10.2307/1931034

33. Donnelly, K. 1978. Simulation to determine the variance and edge effect of total nearest neighbour distance. In Simulation methods in archaeology. Edited by I.R. Hodder. Cambridge University Press, London. pp. 91–95.

34. Ezquerra-Romano, I., & Ezquerra, A. (2017). Highway to thermosensation: A traced review, from the proteins to the brain. Reviews in the Neurosciences, 28(1), 45–57. 10.1515/revneuro-2016-0039

35. Dhaka A., Earley T. J., Watson J. & Patapoutian A. (2008). Visualizing cold spots: TRPM8-expressing sensory neurons and their projections. Journal of Neuroscience, 28(3), 566–575. 10.1523/JNEUROSCI.3976-07.2008

36. Hensel H. & Zotterman Y. (1951). The response of the cold receptors to constant cooling. Acta physiologica Scandinavica, 22(2-3), 96-105. 10.1111/j.1748-1716.1951.tb00758.x

37. Leone, Caterinaa; Dufour, Andréb; Di Stefano, Giuliaa; Fasolino, Alessandraa; Di Lionardo, Andreaa; La Cesa, Silviaa; Galosi, Eleonoraa; Valeriani, Massimiliano; Nolano, Mariae; Cruccu, Giorgioa & Truini, Andreaa (2019). Cooling the skin for assessing small-fibre function. Pain 160(9): 1967–1975. 10.1097/j.pain.0000000000001584

38. Belmonte, C., & Viana, F. (2008). Molecular and cellular limits to somatosensory specificity. Molecular pain, 4(1), 1–17. 10.1186/1744-8069-4-14

39. Paricio-Montesinos, R., Schwaller, F., Udhayachandran, A., Rau, F., Walcher, J., Evangelista, R., … & Lewin, G. R. (2020). The sensory coding of warm perception. Neuron, 106(5), 830–841. 10.1016/j.neuron.2020.02.035

40. Zimmermann, K., Hein, A., Hager, U., Kaczmarek, J. S., Turnquist, B. P., Clapham, D. E., & Reeh, P. W. (2009). Phenotyping sensory nerve endings in vitro in the mouse. Nature protocols, 4(2), 174–196. 10.1038/nprot.2008.223

41. Ackerley, R., & Watkins, R. H. (2018). Microneurography as a tool to study the function of individual C-fiber afferents in humans: responses from nociceptors, thermoreceptors, and mechanoreceptors. Journal of neurophysiology, 120(6), 2834–2846. 10.1152/jn.00109.2018

42. Bouvier, V., Roudaut, Y., Osorio, N., Aimonetti, J. M., Ribot-Ciscar, E., Penalba, V., … & Crest, M. (2018). Merkel cells sense cooling with TRPM8 channels. Journal of Investigative Dermatology, 138(4), 946–956. 10.1016/j.jid.2017.11.004

43. Campero, M., Serra, J., Bostock, H., & Ochoa, J. L. (2001). Slowly conducting afferents activated by innocuous low temperature in human skin. The Journal of physiology, 535(3), 855–865. 10.1111/j.1469-7793.2001.t01-1-00855.x

44. Saal, H. P., & Bensmaia, S. J. (2014). Touch is a team effort: Interplay of submodalities in cutaneous sensibility. Trends in Neurosciences, 37(12), 689– 697. 10.1016/j.tins.2014.08.012

45. Jörntell, H., Bengtsson, F., Geborek, P., Spanne, A., Terekhov, A. V., & Hayward, V. (2014). Segregation of tactile input features in neurons of the cuneate nucleus. Neuron, 83(6), 1444–1452. 10.1016/j.neuron.2014.07.038

46. Mountcastle, V. B. (1957). Modality and topographic properties of single neurons of cat’s somatic sensory cortex. Journal of neurophysiology, 20(4), 408–434. 10.1152/jn.1957.20.4.408

47. Ochoa, J. L. (2010). Intraneural microstimulation in humans. Neuroscience letters, 470(3), 162. 10.1016%2Fj.neulet.2009.10.007

48. Courtin, A. S., Delvaux, A., Dufour, A., & Mouraux, A. (2023). Spatial summation of cold and warm detection: evidence for increased precision when brisk stimuli are delivered over larger area. Neuroscience Letters, 137050. 10.1016/j.neulet.2023.137050

49. Fardo, F., Beck, B., Allen, M., & Finnerup, N. B. (2020). Beyond labeled lines: A population coding account of the thermal grill illusion. Neuroscience & Biobehavioral Reviews, 108, 472–479. 10.1016/j.neubiorev.2019.11.017

50. Ho H.N., Watanabe J., Ando H., & Kashino M. (2011). Mechanisms Underlying Referral of Thermal Sensations to Sites of Tactile Stimulation. Journal of Neuroscience, 31(1), 208–213. 10.1523/JNEUROSCI.2640-10.2011

51. Cataldo A., Ferrè E.R., Di Pellegrino G. & Haggard P. (2016). Thermal referral: Evidence for a thermoceptive uniformity illusion without touch. Scientific Reports, 6(September), 1–10. 10.1038/srep35286

52. Johnson, K. O., Darian-Smith, I., LaMotte, C., Johnson, B., & Oldfield, S. (1979). Coding of incremental changes in skin temperature by a population of warm fibers in the monkey: correlation with intensity discrimination in man. Journal of Neurophysiology, 42(5), 1332–1353. 10.1152/jn.1979.42.5.1332

53. Sabesan, R., Schmidt, B. P., Tuten, W. S., & Roorda, A. (2016). The elementary representation of spatial and color vision in the human retina. Science advances, 2(9), e1600797. 10.1126/sciadv.1600797

54. Mancini, F., Bauleo, A., Cole, J., Lui, F., Porro, C. A., Haggard, P., & Iannetti, G. D. (2014). Whole-body mapping of spatial acuity for pain and touch. Annals of neurology, 75(6), 917–924. 10.1002/ana.24179

55. Amir, C., Rose-McCandlish, M., Weger, R., Dildine, T.C., Mischkowski, D., Necka, E.A., Lee, I.S., Wager, T.D., Pine, D.S. and Atlas, L.Y. (2022). Test-retest reliability of an adaptive thermal pain calibration procedure in healthy volunteers. The journal of pain, 23(9), 1543–1555. 10.1016/j.jpain.2022.01.011

56. Moloney, N. A., Hall, T. M., & Doody, C. M. (2012). Reliability of thermal quantitative sensory testing: a systematic review. Journal of rehabilitation research and development, 49(2), 191. 10.1682/JRRD.2011.03.0044

57. Fagius, J., & Wahren, L. K. (1981). Variability of sensory threshold determination in clinical use. Journal of the Neurological Sciences, 51(1), 11–27. 10.1016/0022-510X(81)90056-3

58. Marshall, K. L., Clary, R. C., Baba, Y., Orlowsky, R. L., Gerling, G. J., & Lumpkin, E. A. (2016). Touch receptors undergo rapid remodeling in healthy skin. Cell Reports, 17(7), 1719–1727. 10.1016/j.celrep.2016.10.034

59. Ohmi, M. (2016). Application to skin physiology using optical coherence tomography. Laser Therapy, 25(4), 251–258. 10.5978%2Fislsm.16-OR-19

60. Baumgärtner, U., Cruccu, G., Iannetti, G. D., & Treede, R. D. (2005). Laser guns and hot plates. Pain, 116(1), 1–3. 10.1016/j.pain.2005.04.021

61. Pennes, H. H. (1948). Analysis of tissue and arterial blood temperatures in the resting human forearm. Journal of applied physiology, 1(2), 93–122. 10.1152/jappl.1948.1.2.93

